# Analysis of the polar residues located at the head domain of focal adhesion protein vinculin under the presence of the *Shigella* effector IpaA and its possible implications during *in vivo* mechanotransduction

**DOI:** 10.1101/2022.11.23.517744

**Authors:** Benjamin Cocom-Chan, Hamed Khakzad, Cesar Valencia-Gallardo, Guy Tran Van Nhieu

## Abstract

Vinculin is a protein associated to linking adhesion receptors facing the outside of cells and reinforcing them by linking it’s intracellular domain of those receptors or, in the case of Cell-Matrix adhesions, to bind to a first level adaptor protein such as talin. The structural organization of vinculin allows it to bind on one part to specific amphipathic motifs collectively designated as vinculin binding sites (VBS), to a set of different vinculin coactivators or actin regulators, and finally a domain responsible to constantly bind to F-actin in a catch bond manner. However, the ability of vinculin to effectively bind all of those intracellular partners, is highly dependent on its structural organization. Which is critically dependent on its ability to respond to mechanical tension on the molecule itself and not necessarily to its binding capacity to VBSs and complementary activators. This is recognized as the combinatorial model of activation. Nonetheless, Shigella’s IpaA effector protein is able to mimic the conformational changes associated with the ones associated with the mechanical deformation of the molecule. This model of vinculin activation is designated as the non-combinatorial model, as the presence of a single activation-partner is enough to get the same effect. This work is devoted to dig in further to develop the previous work from this lab, as we have been able to characterize the *in vitro* and *in vivo* effects of Shigella’s IpaA-Cterm region as the one responsible for both inducing conformational changes in solution, as well as the formation of super-stable adhesion, associated to maturity markers as VASP and alpha actinin. Additionally the IpaA-Cterm transfection renders those cells with the ability to maintain the adhesion structures stable and even resist the action of actomyosin relaxing molecules. Which renders them as mechanically-independent adhesions. We found that residue substitution at the surface of D1 and D2 interphase, (as well as residues maintaining the D2 domain helical bundles folded), might participate in the maintaining the structural integrity and interdomain interaction during force dependent as revealed by its ability to form protein complexes in vitro and under force-independent settings, as the morphology of cellular adhesions is altered in a way different from the previously reported targeting only the D1-D5 interaction.

## Introduction

The combinatorial model of vinculin activation includes, besides the D1-VBS interaction, the participation of auxiliary molecules as phosphatidylinositol-4,5-bisphosphate (PIP2) before being able to bind to F-actin. It has been proposed that mechanical tension is necessary in order to allow the recruitment of vinculin to cellular adhesion sites and the unveiling of its different binding sites allow vinculin to interact with other proteins. This model is referred to as the combinatorial activation model, as it considers necessary the presence of secondary activators in order to overcome the tight intermolecular interaction between the vinculin Vt and other head domains (H. Chen, Choudhury, and Craig 2006; Carisey and Ballestrem 2011).

The presence of an α-helix motif, vinculin binding site (VBS) is an important feature to consider if a given protein is alleged to be considered a vinculin binding partner or activator (Gingras et al. 2005). Nonetheless, the intracellular pathogens such *Shigella* spp. requires the effector protein IpaA, an effector protein with the potential to interact with vinculin by means of its three VBSs already exposed independently of mechanotransduction to cause infection (Park et al. 2011; Kluger et al. 2020).

In terms of biological functions, the most probable conformational (thermodynamic) pathways that a protein is likely to transit between two different states (i.e inactive/active or unbound/bound states) can be recapitulated by analyzing their intrinsic conformational predisposition(s) in its apo/native form (Berendsen and Hayward 2000). Experimental data have showed agreement with this proposal and some researchers have even proposed that this structure-encoded properties imply that: i) a protein posses a fingerprint or unique landscape of dynamic fluctuations (at equilibrium conditions) that depends in its 3D structure; and ii) those fluctuations or predispositions might be functionally or even evolutionary constrained (Tobi and Bahar 2005). Different teams have studied vinculin through different structure-based modeling (SBM) approaches in order to understand the transition between the inactive to active states normally including:

1. the helical bundling conversion after VBS binding/insertion to the D1 domain,
2. the relationship (contacts) among subdomains in the apo/native state and during the activation transitions and,
3. the inclusion of pulling forces as expected after Vt-F-actin interaction in a different direction (or even opposite) from the one acting on the D1-VBS complex.

Nonetheless, the potential role of the vinculin head domain (D1-D4) as an intrinsic regulator during vinculin activation under mechanical tension-responses has gained attention more recently, thanks to different structural and biochemical approaches, unveiling an aspect on vinculin, yet to be clarified.

Overall, *Shigella’s* IpaA ability to induce vinculin activation in the absence of pulling forces, remains a topic to be fully explored. Recent structural models propose major conformational reorganizations on vinculin while preserving its folded integrity, a property related to the protein architecture and its ability to resist unfolding under mechanical tension (Forman and Clarke 2007). Examples of this mechanical stability have been identified in spring-like ankyrin domains (Lee et al. 2006) and as it has been suggested by biochemical data and MD predictions (Le, Yu, and Yan 2019; Chorev et al. 2018; Kluger et al. 2020). Recently a model of activation has been proposed involving a reorientation of all the four pairs of helical bundles from vinculin, while D5 (Vt) does not separate from the other domains (Stec and Stec 2022).

Vinculin represents a challenge regarding the dissection of the relationship between its possible conformational states (depending on which molecule(s) vinculin is interacting with) and the function(s) associated with those conformational states. In order to understand the initial conformational states of vinculin activation, IpaA C-term can be used as a tool to better understand the molecular mechanisms leading to vinculin activation, as it possesses a unique capacity of permitting vinculin to acquire its active conformation and bind to F-actin in the absence of additional signals, (compared to the >2 input signals model of activation required normally in cells) (H. Chen, Choudhury, and Craig 2006).

As such, *Shigella*’s IpaA domain possesses three different VBS located in it’s C-terminus domain, and it represents an excellent tool given its ability to mimic the properties of (otherwise) mechanically sensitive events occurring in cells under mechanical stress) in vitro, with an ability to bind at very low Kd (pM range) than their endogenous VBS partners.

Differences should be considered, between the *Shigella’s* IpaA dependent vinculin activation and the activation of vinculin under mechanical forces. As its name indicates, mechanical unfolding involves the action of pulling forces in a protein. The main difference resides in the conformational pathway(s) the same protein can transit between two different configurations depending on the nature of the stimulus (either chemical, mechanical or thermal unfolding). Not every protein can maintain its structural integrity under mechanical stress (Stirnemann et al. 2014), as such, the mechanical sensitive proteins represent a special case when its structure is tightly related to its function. They have a specific architecture that maintains their inter and intramolecular interactions such as the non-covalent bonds or polar interactions, while resisting the mechanical tension without unfolding of the local or domain level structures even after major conformational rearrangements occur (Y. Chen, Radford, and Brockwell 2015). This aspect is crucial in order to properly translate a mechanical input into a biochemical and biological response (i.e. unveiling a signaling or binding domain or a post translational modification site).

Previous SEC-MALS analysis of D1D2:IpaA-VBSs protein complexes, showed the presence of different protein complexes when mixing purified vinculin derived D1-D4 or D1D2 subunits with the IpaA-VBS_123_. Moreover, after transfecting cells with IpaA-VBS_123_ rendered them with the ability to form stable adhesions resistant to focal adhesion disassembly even after applying acto-myosin relaxing molecules Y27632 (Valencia-Gallardo et al. 2022). The protein complexes corresponding to the 1:1 molar ratio were isolated and structural models were obtained by TX-MS structural modeling (Valencia-Gallardo et al. 2022).

As the molecular mechanism driving the allosteric changes observed in vinculin D1D2 subunits formation after IpaA-VBS_123_ protein-protein interaction remained elusive, the aim of this study is to characterize the molecular mechanism driving the IpaA-dependent vinculin activation in the absence of mechanotransduction and its possible role in vivo. First, we classified the rate of in vitro monomer to oligomer conversion of D1D2 after the interaction with IpaA-VBS123 and the composition for the different bands observed in Native PAGE gels. As control experiments, we compared the D1D2 monomer to oligomer conversion efficiency by comparing the conversion rate after interaction with IpaA-VBS123 wt and a variant version at residue 499 located in the VBS3, the contact region found to be interacting with the D1DD2 cleft accordingly to the TX-MS structural model. After inverting the charge from Lysine to Glutamate, we found a reduced capacity of higher oligomer formation, accompanied with slight intensity increase for the bands corresponding to the intermediate complexes in native PAGE conditions compared to the WT protein. This suggests that a residue charge inversion located in VBS3 reduces its capacity to form higher D1D2 complexes. However, as VBS3 can bind to D1 as described in its original characterization (Park et al. 2011) (the same way as VBS1) and the D1D2 cleft region (as the TX-MS models suggests) it isn’t clear if this deficiency might correspond to which binding site or both (Figure 1A)

**Figure 1.**
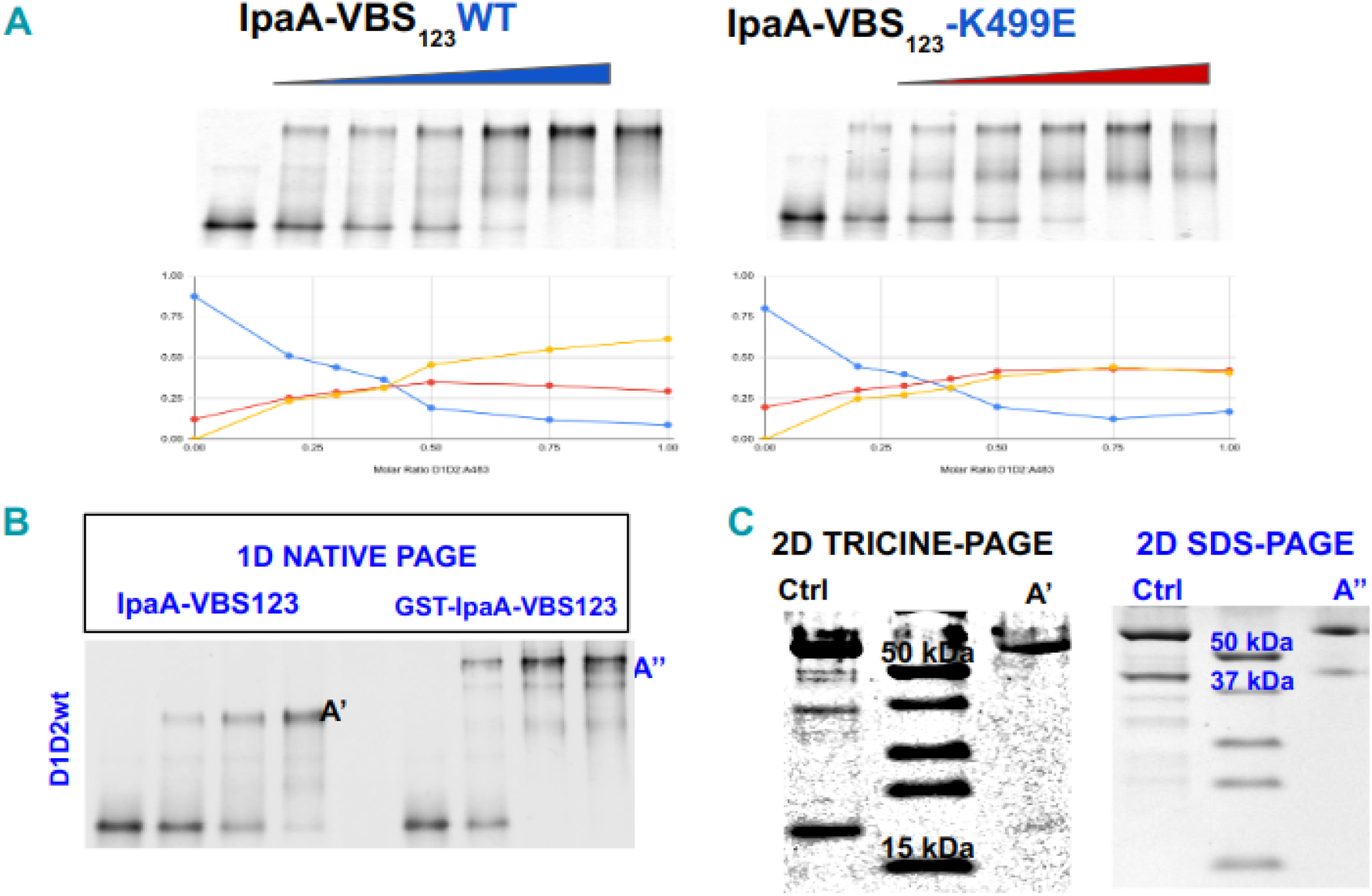
IpaA-Cterm form different protein complexes with the vinculin head domain subunits D1D2 and structural models suggest a role for VBS3 a conformational change on D2 relative to D1. **A)** In native PAGE gels experiments ran after in vitro protein:protein interaction complex formation by titration of vinculin D1D2 with IpaA-VBS123 showed the appearance of different complexes with a prominent band corresponding to the higher oligomeric form. To the right are presented the same complexes formed with different abundances when introducing a charge inversion VBS3 (IpaA-VBS123-E499K). **B)** The higher oligomer complexes remained stable during native gel migration after incubation with the cleaved and non cleaved (GST-fusion) of IpaA-VBSs, and the major bands derived from NATIVE-PAGE gels can be quantified and further analyzed in a second dimension SDS-PAGE gel. **C)** Bands corresponding to vD1D2 and IpaA-Cterm are detectable.

**Figure 2.**
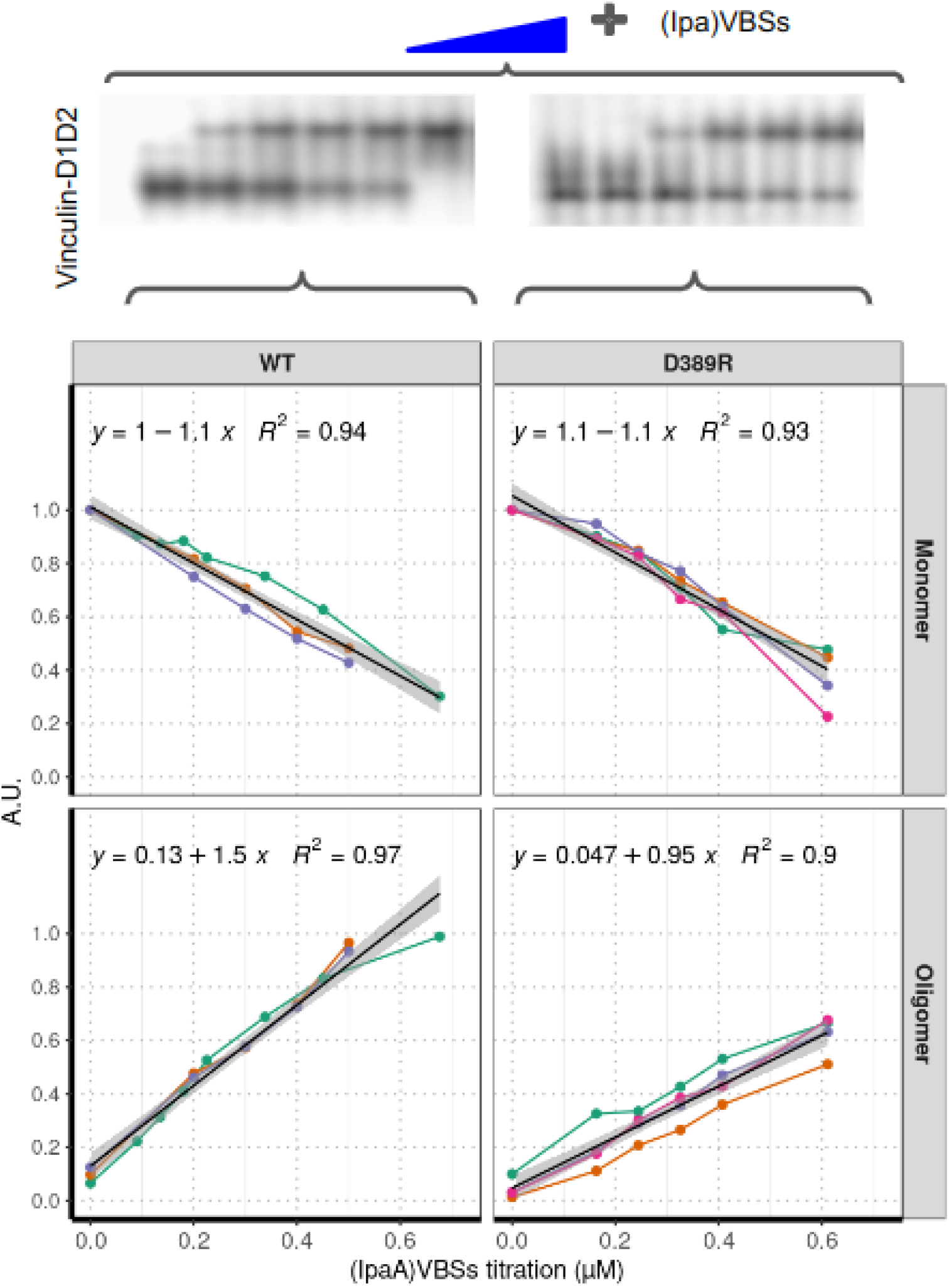
Vinculin point variants revealed a role for the IpaA-VBS_123_ to induce the formation of stable complexes at different titration rates compared to the D1D2 wt version. A) Comparison for gels quantified in order to show the rates of monomer disappearance and oligomer formation as a measurement for the efficiency of interaction between D1D2 with IpaA-VBS123 between the WT and and the D389R version. Independent experiments are shown in different line colors.

## MATERIALS AND METHODS

### Materials and Methods

List of sequences for primers used for the site directed substitution of the human vinculin D1D2 construct cloned in a pET15b and the pGex-4T2-IpaA-VBS123 is the following:

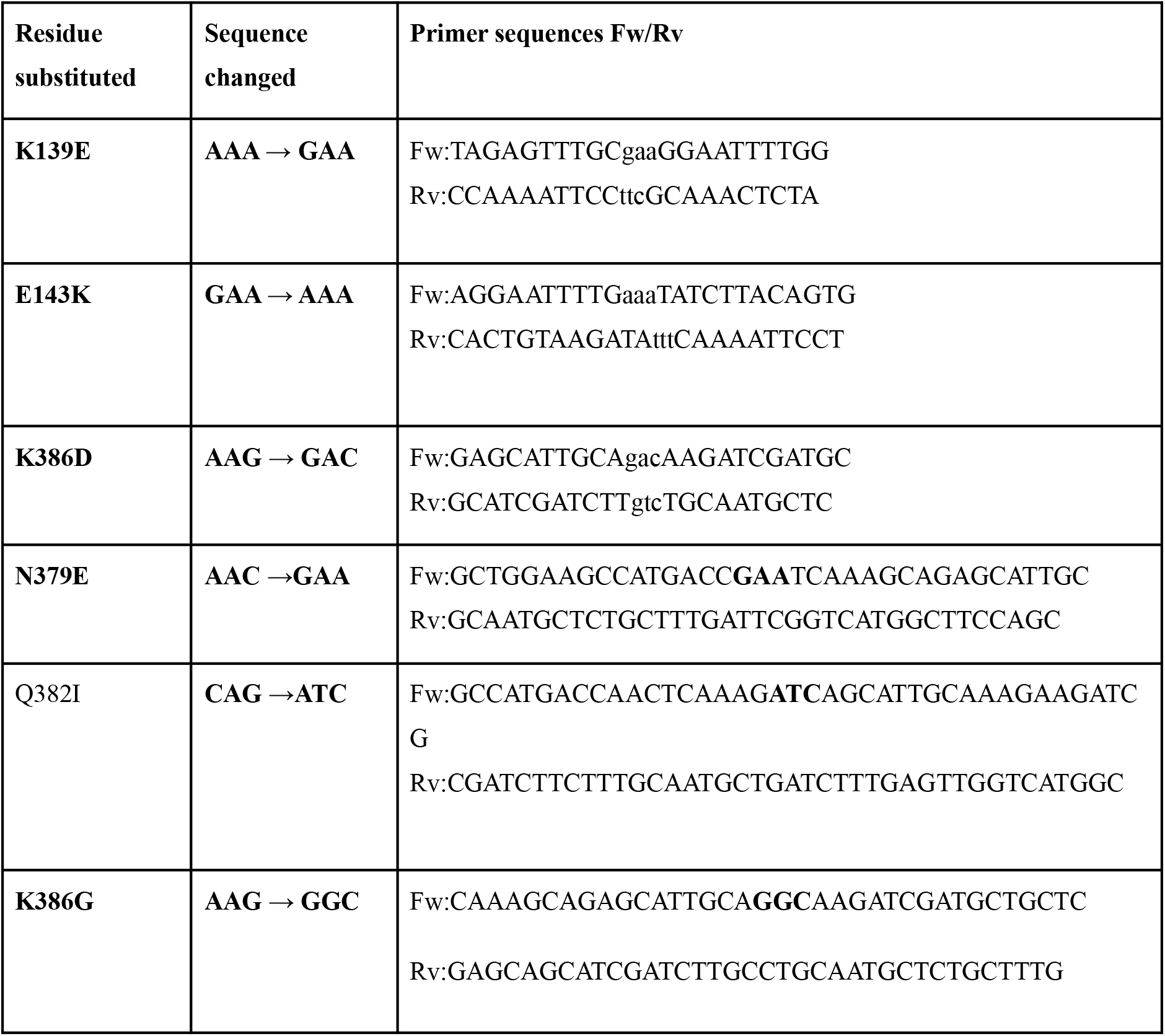
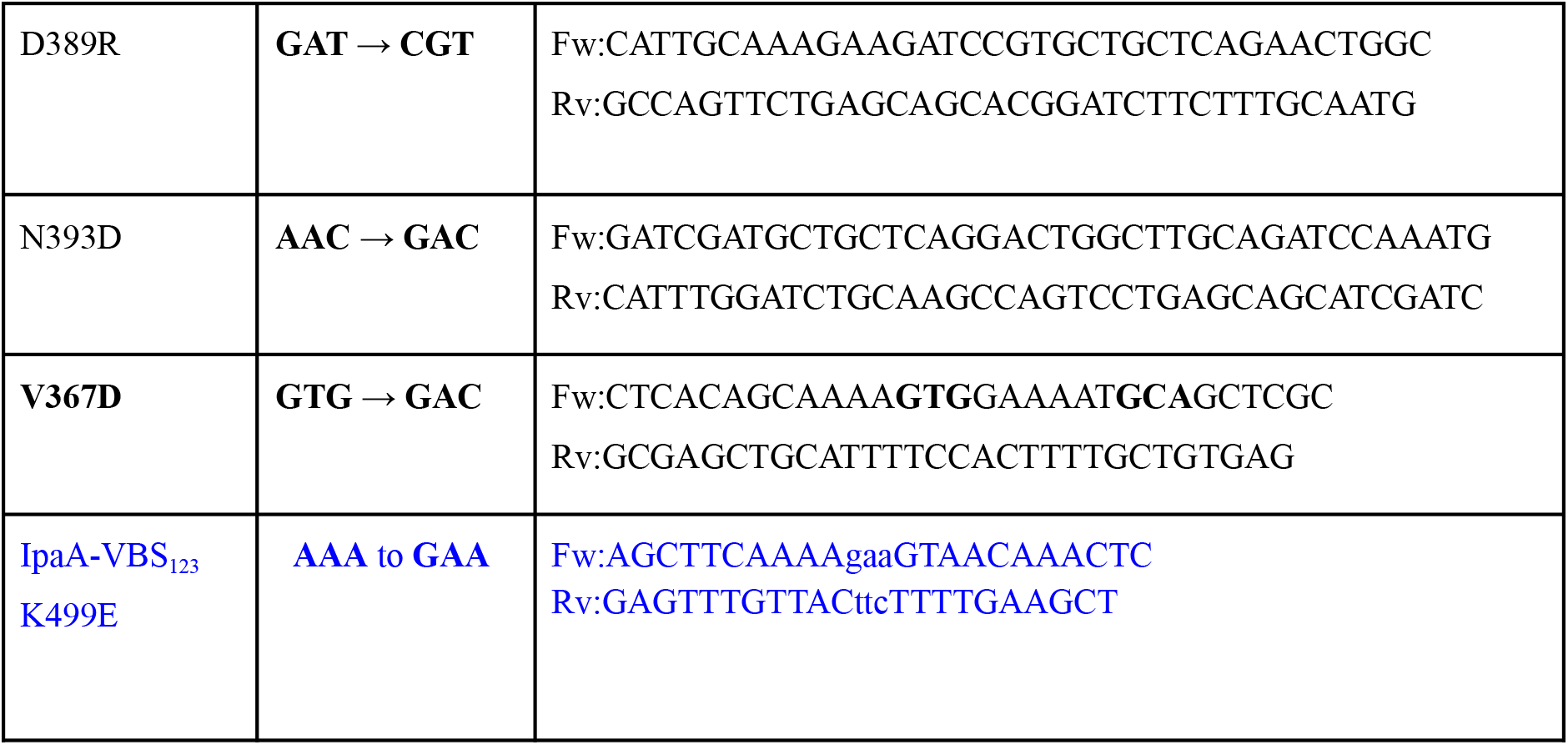

### PCR Reaction and Program

#### 50 μl per reaction/tube

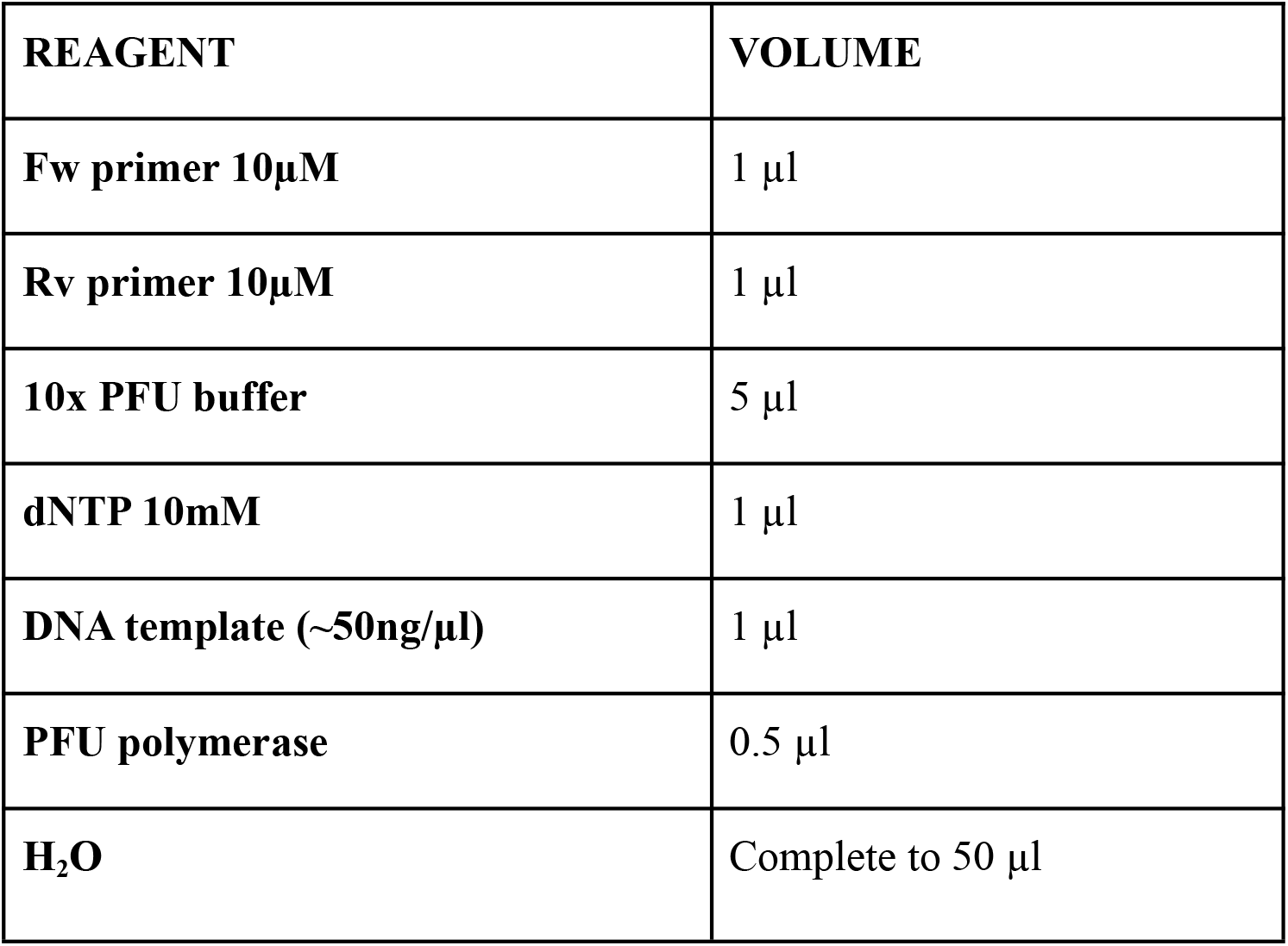

The amplification product was treated for a DPNI matrix (1U) digestion for 90 min at 37 °C and 80 °C for 20 min 4 °C pause. This reaction was used to transform chemically or electro-competent bacteria for subsequent cloning and sequencing and protein purification.

The regions flanking the residues mutated in the hvD1D2-pET15b constructs were double digested with PspXI and BclI restrictions enzymes and further transferred using the T4 ligase (according to New England Biolabs online instructions) to full length Vinculin-mCherry-N1 constructs previously treated with the same enzymes. The effectiveness for transference was corroborated by Sanger sequencing.

### Protein Purification

Custom prepared BL21 chemically competent E. coli cells were transformed with either pet15b-vhD1D2 or pGex-4T2-IpaA-VBS-123 plasmids (wt or variants). After reaching an O.D. of ∼1 at 600nm, the culture was induced with IPTG for 3 hours before bacteria was lysate in a microfluidizer or french press. The D1D2 was purified as described in Valencia-Gallardo et. al by using a lysis buffer with 25mM Tris–HCl pH 7.4, 0.5M NaCl (including protein inhibitors accordingly Complete or Pefabloc) and 30mM imidazole using ∼5 ml Niquel beads and eluted at 20mM Tris-HCl (pH 7.4), 0.5M NaCl, 300 mM Imidazole (pH 8.0). For the GST-IpaA-VBS-123 lysates were dialyzed in 25 mM Tris–HCl pH 7.4, 0.5 M NaCl and 30 mM imidazole and captured with Glutathione-Sepharose 4B beads as indicated by the manufacturer (GE Healthcare). The elution fractions were pooled and the buffer was exchanged to phosphate-buffered saline (PBS) and digested with thrombin followed by ion exchange chromatography column and concentrated with a 3 kDa cut-off filtration unit (Amicon). Protein concentration was determined using the BCA assay (Thermo Scientific). Samples were dialysed in 20mM Tris-HCL (pH 7.4), 100mM NaCl, 1mM β-Mercaptoethanol for the protein interaction experiments in vitro and clear native PAGE gels.

## NATIVE PAGE PROTOCOL

### IN VITRO REACTIONS

The D1D2:IpaA-VBS123 in vitro protein:protein interactions were performed in 20mM Tris-HCL (pH 7.4), 100mM NaCl, 1mM β-Mercaptoethanol buffer utilizing fixed concentrations of D1D2 (∼10 μM in 20 μl final vol. reaction) at increased titration with IpaA-VBS123 for 30-40 min at RT before mixing with 20 μl native loading buffer (working concentrations of 62.5 mM Tris-HCl, pH 6.8 with 25% glycerol and 0.01% Bromophenol blue).

### RUNNING CONDITIONS FOR NATIVE GEL SHIFTS

Aliquots of 10/40 μl were loaded into 7.5% polyacrylamide mini-gels (1mm width) by using standard gel recipes but substituting SDS for water. Runnings were performed during 150-180 min at 25mA per gel were used in order to observe the bands corresponding to the monomers and the higher order oligomers 3:1 or oligomer (cold conditions were preferred during the runnings). D1D2-E143K was the only one requiring 180 min in order to observe higher bands. Molar ratios of D1D2:IpaA-VBS123 were between 1:0→ 1:1. The gels were stained using colloidal Coomassie R-250 blue.

### GEL IMAGING AND QUANTIFICATION

Band intensities corresponding to the monomer, higher order oligomer or intermediate species were measured using ImageJ with a fixed rectangular area (adjusted with the max width for the monomer and max high order oligomer for the height). A reference area outside the lanes used for the running reactions was included to subtract the background. Values were plotted and then normalized in order to compare the in vitro phenotypes between D1D2 wt and the different variants. Additionally we included the gell images

### CELL TRANSFECTION EXPERIMENTS

40K/well mouse endothelial fibroblast vinculin null cells (MEF vinc -/-) cells were seeded in 12 well plates 24 hours before transfection. Acid washed coverslips were covered with 0.5 ml of fibronectin 20μg/ml and mounted in the plates before seeding cells. DMEM media with high glucose content (4.5 mg/ml) supplemented with SFB 10% and non essential amino acids (no antibiotics were used.

Transfection using the hFLVmCherry-N1 constructs for the WT and the variant versions transferred using the Fugene reagent and following the vendors instructions. Per well reactions used 1.6 μg of DNA and 3X Fugene reagent diluted in OPTIMEM media to a final volume of 100 μl/ml. The coverslips were treated for 20 minutes with PFA 4%, and processed for fluorescence microscopy, and mounted on slides using Dako mounting medium (Dako, Agilent Technologies), as described (Romero et al., 2012).

## IMAGE ANALYSIS FOR THE FOCAL ADHESION QUANTIFICATION

For Cell Adhesion quantification was performed using ImageJ-FIJI adapting a protocol previously described Horzum et al. 2014 (Horzum, Ozdil, and Pesen-Okvur 2014):

## PLOTS AND GRAPHS

The graphs corresponding to the main figures were plotted using R (https://www.r-project.org/) and standard libraries (Villanueva and Chen 2019).

## RESULTS

The first goal of this study was to evaluate the role of IpaA-VBS3 in the monomer to oligomer conversion of D1D2 after its interaction with IpaA-VBS_123_ wild type or a version with a charge inversion located in the third VBS (IpaA-VBS_123_ -K499E). We compared the D1D2 monomer to oligomer conversion efficiency by comparing the rate after interaction with IpaA-VBS123 wt and a variant version at residue 499 located in the VBS3, a contact region predicted to be interacting with the contact region between D1 and D2 accordingly to the previously identified structural models corresponding to 1:1 hetero-complex (Valencia-Gallardo et al. 2022).

After inverting the charge from Lysine to Glutamate, we found a reduced capacity of higher oligomer formation, accompanied with slight intensity increase for the bands corresponding to the intermediate complexes in native PAGE conditions compared to the WT protein. This suggests that a residue charge inversion located in VBS3 reduces its capacity to form oligomers with D1D2 (Figure 1A). However, as VBS3 can bind to D1, and it was proposed that IpaA-VBS123 can bind simultaneously to three D1 proteins (Park et al. 2011), we isolated the bands corresponding to the higher oligomeric form detected in native PAGE gels conditions (bands A’ and A’’ in Figure 1B) and performed a second dimension gel under denaturing conditions. We observed bands corresponding to both cleaved and uncleaved protein variants containing the three VBS: IpaA-VBS123 and GST-IpaA-VBS123 in second dimension denaturing gels, suggesting the stability of those complexes during the gel migration (see Figure 1B and 1C).

**Table 1.**
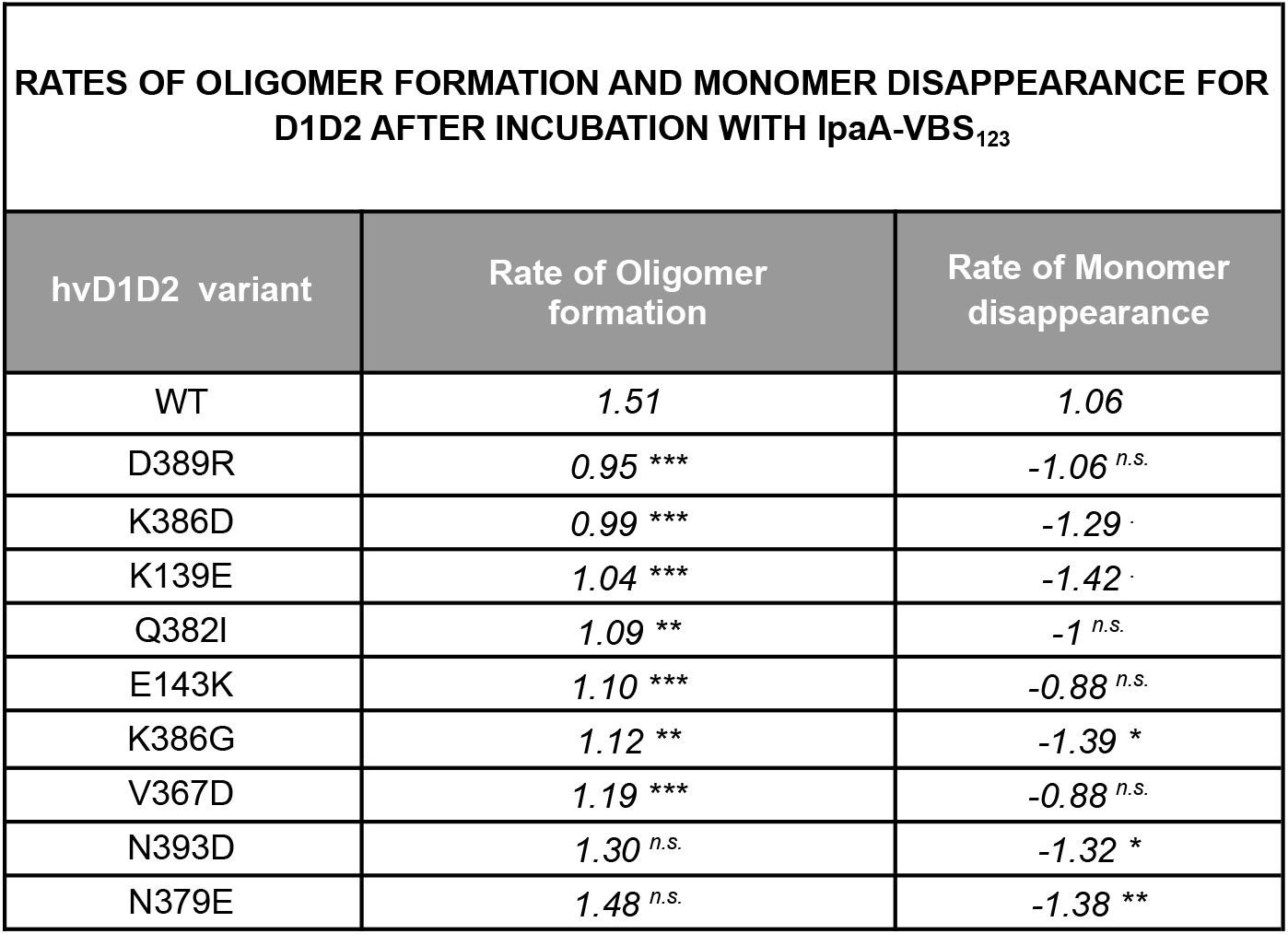
The same slope regression analysis showing the exact values measured in figure 2B, indicating its statistical significance as described in the same figure. Slope values were compared to WT with ANCOVA test Signif. codes: 0=‘***’, 0.001=‘**’, 0.01=‘*’, 0.05=‘.’, 0.1=‘n.s’.

In general, in this section it was possible to characterize the ability of Shigella’s IpaA-VBS123. To form stable oligomeric complexes in vitro but the efficiency in oligomer formation was dependent on the ability of VBS3 to efficiently interact with the D1D2 residues located at the contact interface between both subdomains. This might be relevant in terms of the ability of the protein to respond to mechanical stimuli in the context of the full length vinculin protein and evaluating its properties in vivo was an important next step to take into account.

## CELL BIOLOGY

The phenotype associated with the vinculin loss of function mammalian cells (MEF vinc -/-) disrupts the initial formation of multiprotein complexes, the efficiency of multiple protein-protein interactions, maturation and growth of adhesions (ref). In the longer range those molecular effects can affect cellular aspects such as a reduction in stress fibers, lamellipodia extension and ultimately cell spreading, the associated effects ultimately affect mice development (Xu, Baribault, and Adamson 1998).

Regarding the cell biology, is important to remark that we measured the morphology of cell adhesions based on the most prominent features identified by a in an exercise to making an overview of the microscopy images, so this analysis is mostly based on the relationship between the central and peripheral cell adhesions as a global way to interpret the possible phenotype occurring in the cell in terms of intracellular tension.

In general by residue substitution on the D1D2 sites located in the head domain different from the VBS binding sites or the Vh-D5 contact residues, we observed a reduction in the ability for cells to form adhesions as compared to the vinculin WT transfectants Figure 3. The cell spreading area was also significantly reduced in those cells which suggested a general impact at cell level of functions.

**Figure 3.**
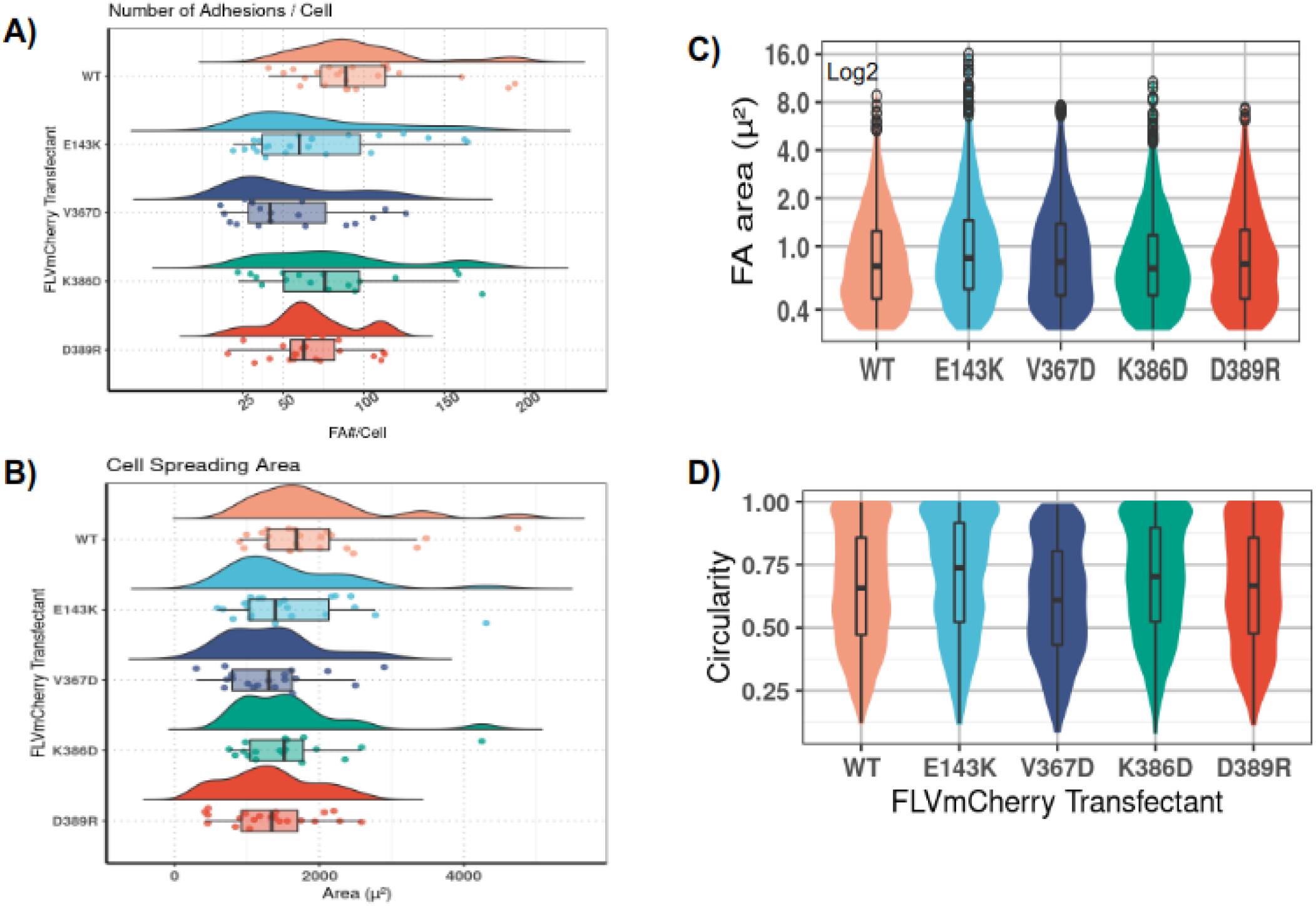
Vinculin -/- MEF cells variants showed transfected with different hFLVmCherry protein variants. The general phenotype observed in cell transfectants was a reduction in cell spreading area A) and less capacity to form adhesions compared to the wt version as indicated in panel B) Additionally, the quantification analysis for the individual adhesions showed differences in terms of size (μm^2^) C) and importantly on circularity index D) the more characteristic shape measurements corresponded to the E143K (comparatively more rounded) and V3667D (more elongated). The dots indicated for the violin plots in panel C) correspond to the outliers.

**Figure 4.**
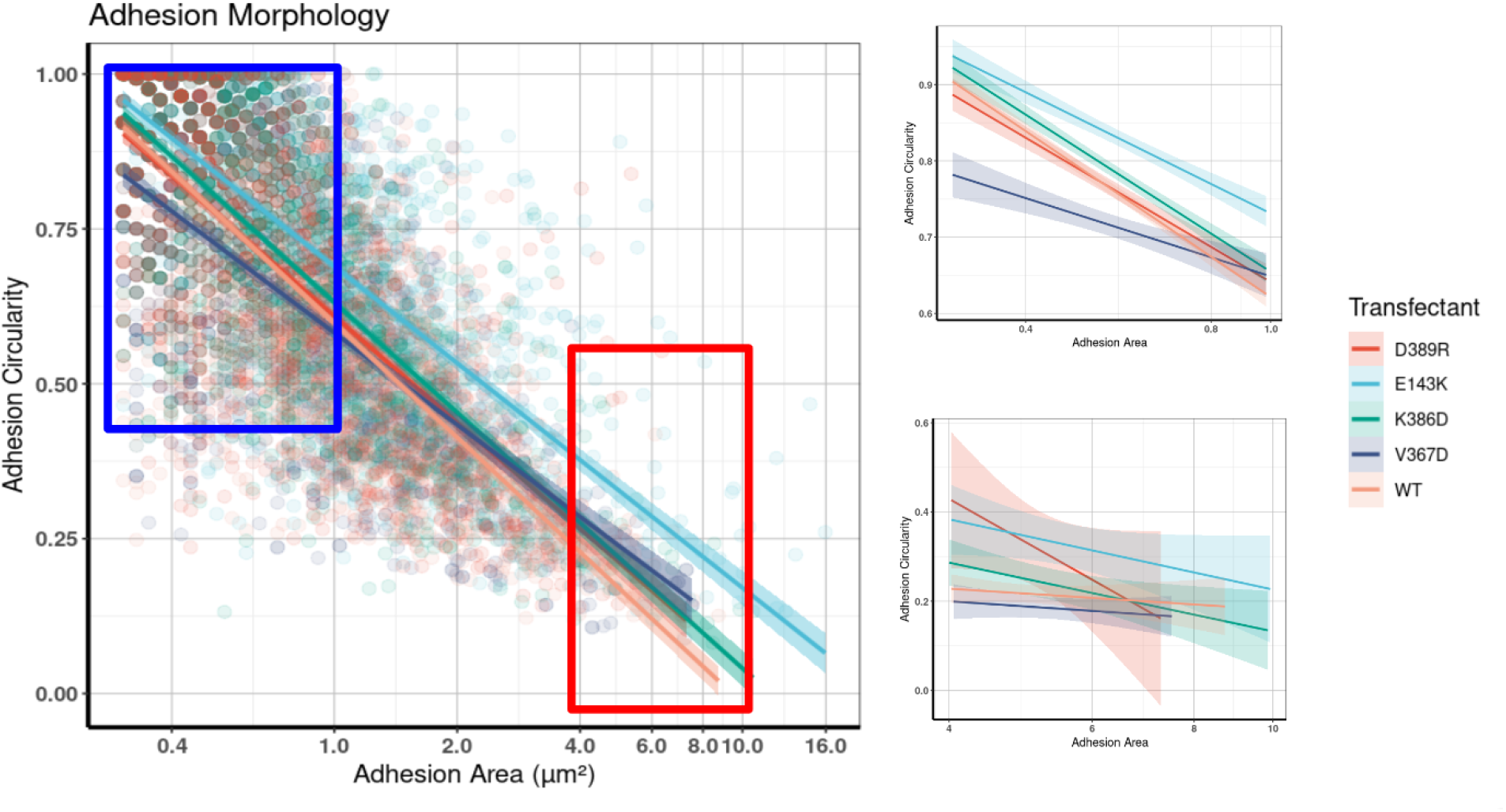
A) Focal adhesion size and shape share a relationship that might resemble the connection of the tail domain to the actin-cytoskeleton. variant versions of hFLV show different values that might resemble its ability to couple properly the connection between the tensile forces to other adhesion cell adhesion proteins like talin, and the tail domain with F-actin. B) Interestingly, the differences observed in this area to circularity and more evident for the smaller adhesions were V367D and E143K showed a trend to form more elongated or rounded adhesions respectively.

Regarding the adhesions per se, we need to consider that any phenotype observed corresponds to a crowded environment but the main difference in phenotype should be attributed to vinculin, and in this case, to a single amino acid substitution. Considering this scenario, resulted evident that the effect on focal adhesion morphology was more considerable altered in terms of size and shape (Figure 3C and 3D).

## DISCUSSION

The possible implications of the different protein complexes observed in the native PAGE gels should be further addressed in terms of determining the stability of those complexes and possibly its lifetime. As the binding of the VBS1 and VBS2 to the vinculin D1 domain has been determined to be at the nM and pM range, this might explain why those complexes remain stable during its migration during native PAGE conditions. The identification of IpaA-VBS123 in second dimension gels under denaturing conditions was supported by the recovery of bands corresponding to IpaA-C-term derivatives in its cleaved IpaA-VBS123 and noncleaved form GST-IpaA-VBS123. Suggesting that the 2:1 and 3:1 complexes are the predominant oligomer forms. The relative proportion between the different complexes in the mixture remains to be determined and this might probably lead to better the succession of conformational changes that the head domain is experiencing during mechano-transduction.

The ability of IpaA to induce vinculin activation in the absence of mechanotransduction suggests that IpaA-VBS123 can use a mechanism at least similar to the one occurring in cells in terms of conformational changes induced under force transmission events. Identifying the molecular player accomplishing this role in cells remains to be covered in the future, however as there exists aminoacid sequence similarities between the IpaA-VBS3 and the helix H46 from talin’s and both can stably bind to talin’s H1-H4 protein derivative. This suggests that H46 might be responsible for the same role as IpaA-VBS3 in terms of interacting with a region different from vinculin’s D1 region, and induce the conformational changes associated with force transmission events. Further experiments such as AFM with purified H1H4 and D1D2 proteins to corroborate this hypothesis are needed, as the case of the talin rod domains needs the exertion of mechanical pulling forces in order to be fully extended and the unveiling of the VBS occurs in comparison to the IpaA-VBS123.

A result not particularly remarked in the results section is that metavinculin might include an endogenous vinculin binding site not reported before. This might have important implications in vinculin biology research to be fully explored ahead (data not showed).

Evidence for the in vivo structure and functions D1D2 and full length vinculin oligomeric forms should be addressed in the future. As we observed that the D1D2:IpaA-VBS123 3:1 hetero-complex, to be stable even under the native gel running conditions, this particular stoichiometry might have implications in vivo to be fully elucidated. However, this needs to be corroborated and more experiments are needed in order to address its possible roles in vivo. One possibility is to act as an actin bundling protein at distance, probably as a short term molecular memory for the pulling direction being exerted from the acto-myosin cytoskeleton associated to the cell adhesion sites, a role that in vivo occurs for example during mechanical-pulling dependent calpain cleavage of the talin rod domain, a function that has important implications in vivo during cell adhesion growth (Saxena et al. 2017).

IpaA might exploit an alternative pathway of activation as the one used by Shigella during infection than the one found in vivo: as such, IpaA has been reported to bind to the D1 domain as other VBSs do by binding to the first helical bundle in D1. However, the presence of VBS1 and VBS2 allows VBS3 to interact with the interphase between D1 and D2 and induce a conformational change that might drive the rotation of D2 respecting D1 in a counterclockwise manner (Valencia-Gallardo et al. 2022). However, the rotation it suffers under mechanical tension, according to SMD simulations from Kluger et al. indicates that D2 rotates respect to D1 in a clockwise direction. Nonetheless, both models should lead to the same result as they involve a major conformational change in the vinculin head domain in order to induce the release of the tail domain to couple the mechanical tension after associating with F-actin.

Substitution of residues located at the interphase of D1-D3 contact regions might also lead to reinforce the idea that the vinculin head domain is playing an important role in terms of acting as a regulator during mechanically-dependent vinculin conformational changes. Additionally, the D2-D3 interphase might play a similar role in regulating the amount or degree of conformational plasticity during mechanically dependent or independent changes in conformation. Those changes associated to the vinculin head domain reorganization that lead to the exposure of the F-actin binding site in the D5 domain and other potential binding partners, might depend on the collective behavior of the protein. The interdomain region shows a strong stability and their binding interactions, they might be coevolving in order to permit the protein to maintain its proper function.

As our results suggest, a single nucleotide substitution could have an impact on the phenotype at the level of cell adhesion morphology and proper cell spreading. As observed by residue substitutions located at the surface of the D1 or D2 domain distant from the VBS-interacting region with VBS3, or the D2, and even at a residue distant and even buried into the H4-helix but that might be important to keep the integrity of the domain and it’s probable due to its resistance to mechanical deformation.

This is an scenario important to consider as the prediction of the coiled-coil motif with their particular combination of hydrophobic and charged residues is located exactly in the middle of the alpha-helix and connecting the two bundles of D2. This is an indication about the importance of keeping its structure in the folded form in order to resist unfolding under mechanical tension. And as the structural predictions based on the vinculin full length folded form suggests, D2 is one of the subdomains more propense to experience the major conformational changes among the other vinculin subdomains as its axis of orientation respecting to the angle of the pulling force from the VBS bound to D1 is almost perpendicular respect to the other subdomains before the exposure of the actin binding site that allows vinculin to perform its connecting function. This is among the first pieces of evidence that, if further confirmed, might implicate a considering vinculin as a mechanosensor protein and not only an scaffolding or adaptor in cell adhesion structures.

The observation that cells transfected with the full length vinculin protein with the residue substitution V367D present adhesions structures already elongated at the initial steps of formation, might suggest that the protein is more sensitive to suffer conformational changes even at low levels of pulling forces from the actin cytoskeleton and even before the tension from myosin pulling is exerted and the adhesions can fully mature. The implications at the level reduced the efficiency of cell adhesion formation with a concomitant reduction in cell spreading area.

The phenotype corresponding to the mutation at the residue E143K located in D1 showed an opposing effect in terms of adhesion shape, as those cells apparently formed rounder adhesions than the WT along all size ranges but similar to the V367D, they formed less adhesions and spreaded less.

Altogether, the substitution of single residues altering either the proper interactions at the surface of interdomain interactions or that might alter the proper folding of the domain itself, as is the case of the V367D substitution, can have a larger range effects at the level of altering the size and anisotropy of focal adhesions at different stages of maturation, as inferred from the adhesion area, suggests that the contact surface between the domains belonging to the vinculin head are key to maintain its conformational changes responding properly to the mechanical inputs.

Additional cell transfection experiments worth exploring in the future are related to the state of maturation of the adhesions at the level of maturation markers such as VASP and high tension markers by evaluating the recruitment of a-actinin will be essential to test in future experiments. Also, it might be interesting to evaluate the extent of migration capacity of cell transfectants compared to the WT.

As for experiments that might clarify the molecular mechanisms underlying the cell adhesion phenotypes and cell spreading, purifying different full length vinculin variant versions and perform in vitro F-actin bundling and cosedimentation assays might help to clarify about the propensity of the different proteins to become in its activate form. Also the possibility of comparing the extent of resistance to protein unfolding as well as SMD simulations, shouldn’t be discarded to further characterize on the role of vinculin’s head domain as a regulator under mechanotransduction.

## SUPPLEMENTARY INFORMATION

**Fig. S1.**
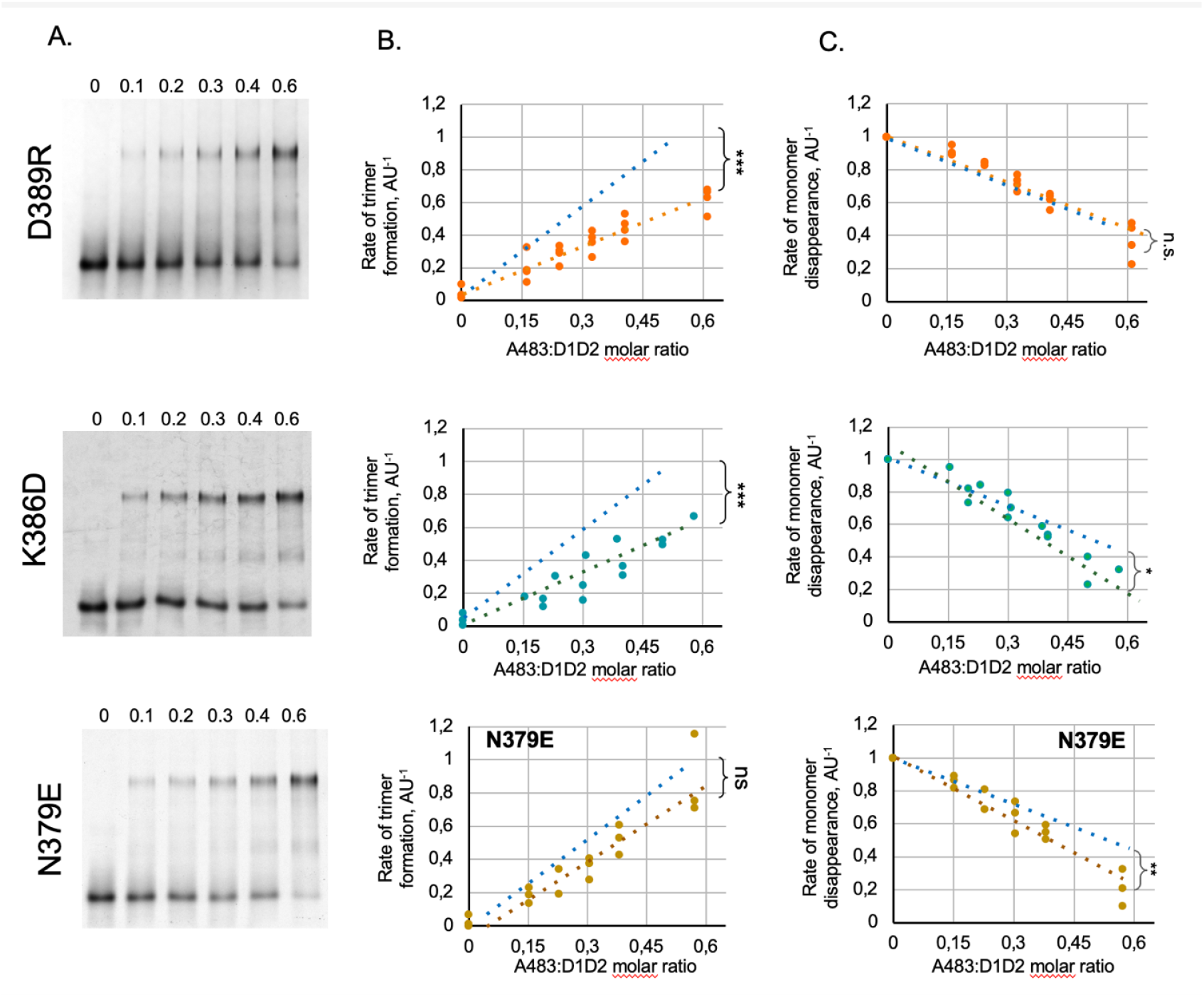
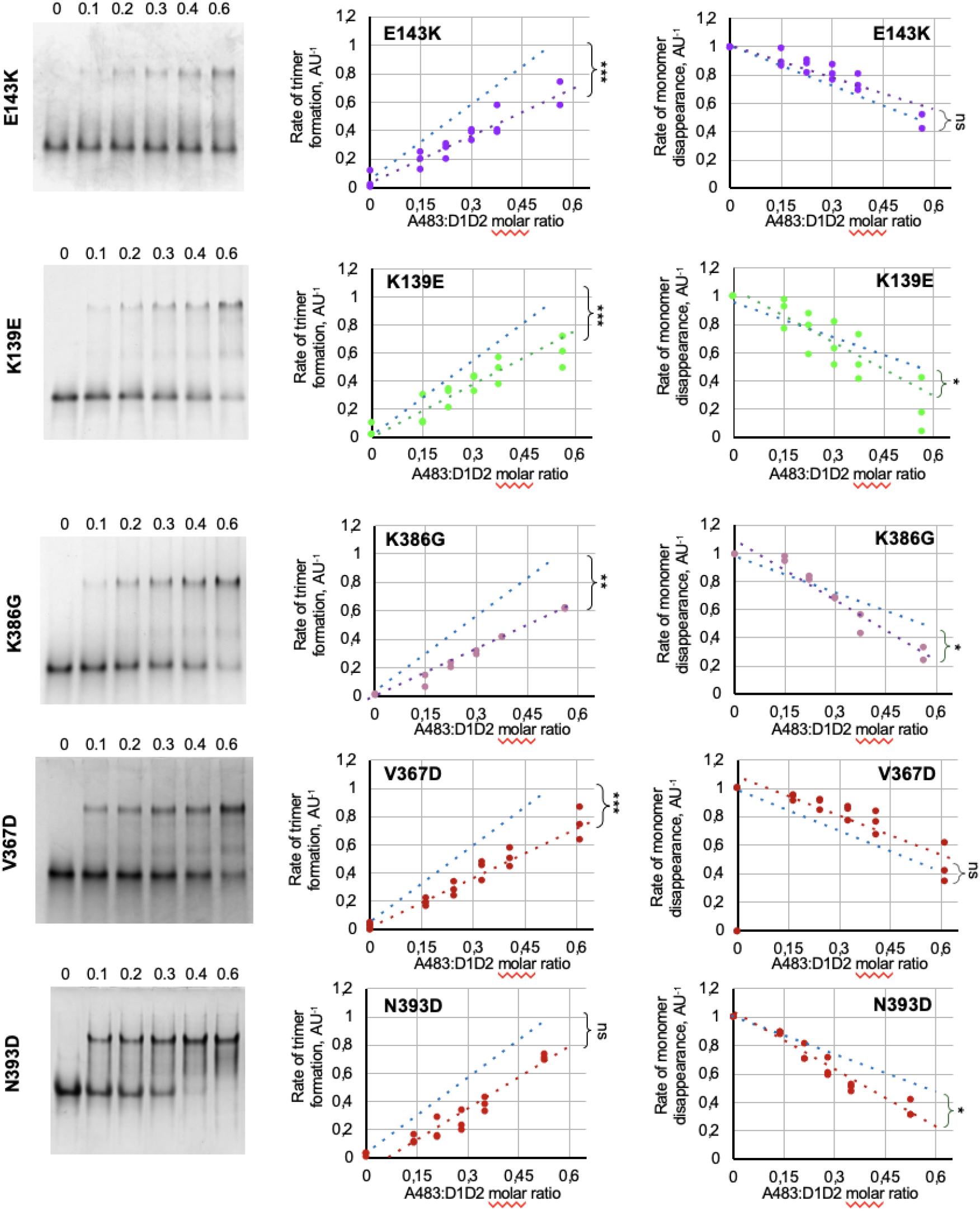
Effects of mutations on D1D2 oligomer formation and monomer disappearance. Representative gel images (A) and quantification corresponding to the rate of oligomer formation (B) and monomer conversion (C).

**Fig. S2.**
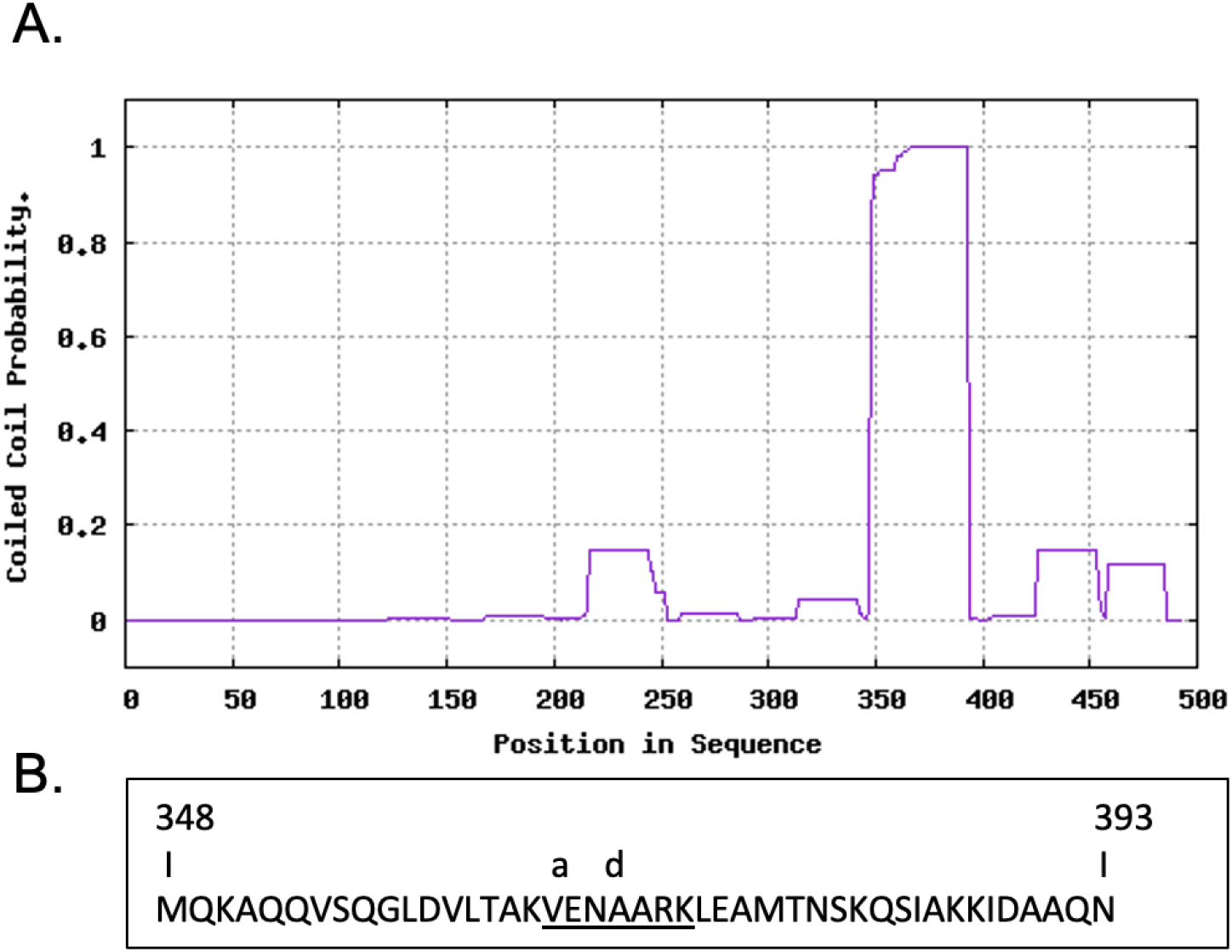
Prediction of a coiled-coil motif in D2 (A), and the sequence targeted in the C-C motif (B) the residue corresponding to the valine was substituted by an aspartate in order to introduce a negative charge and alter the organization of the whole D2 domain.

**Fig. S3.**
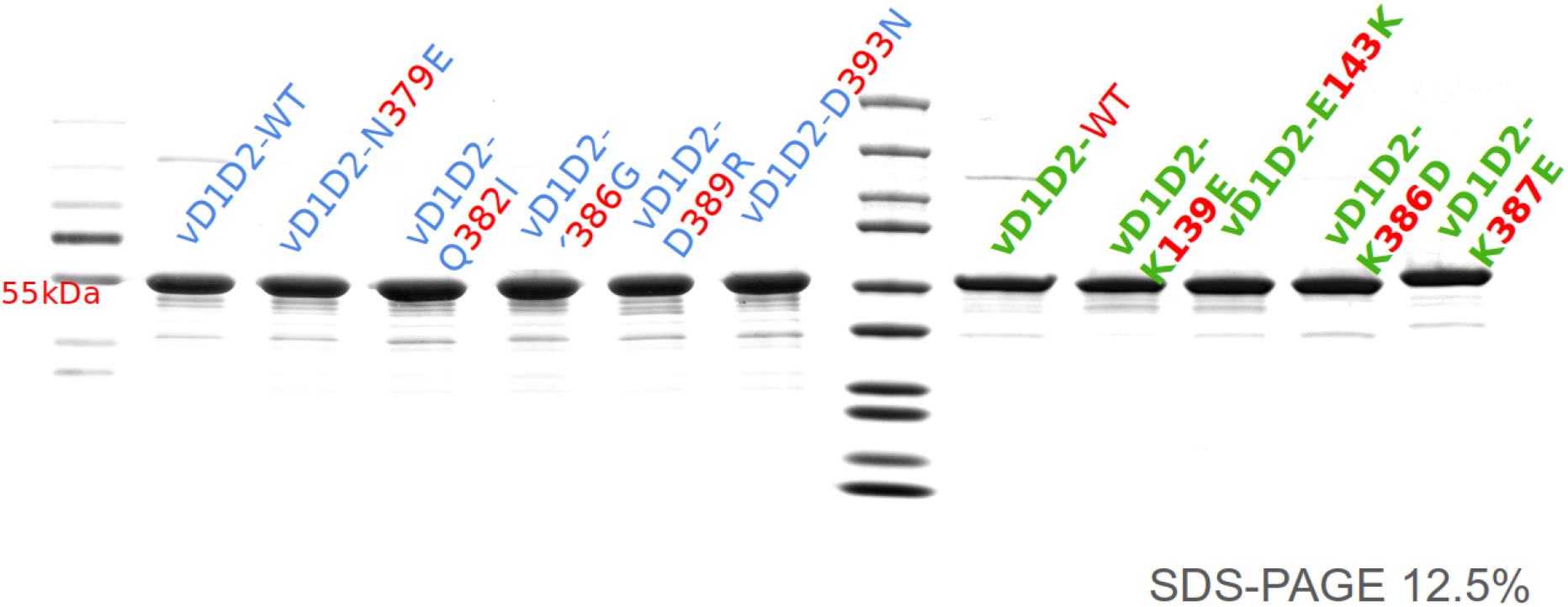
SDS-PAGE corresponding to the D1D2 protein variants used for the in vitro protein:protein interaction experiments. The gel corresponding to the V367D variant is not included in this image as it was purified later.

**Figure S4.**
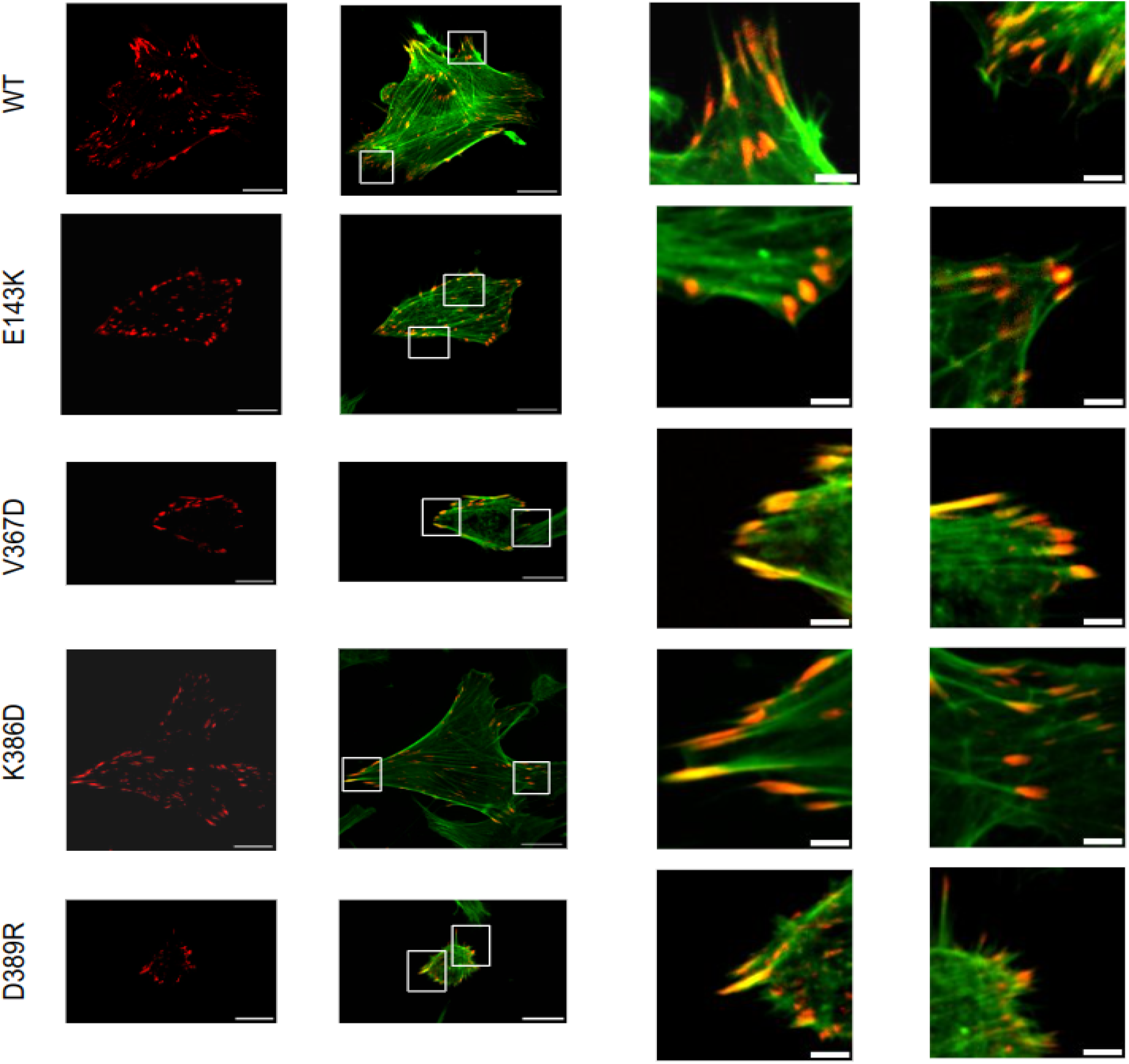
Representative images and insets for the cells transfected with full length vinculin proteins fused to the mCherry fluorescent protein. The variant is indicated on the left side. Scale bars correspond to 20 and 4 μm^2^ respectively.

**Table S4.**
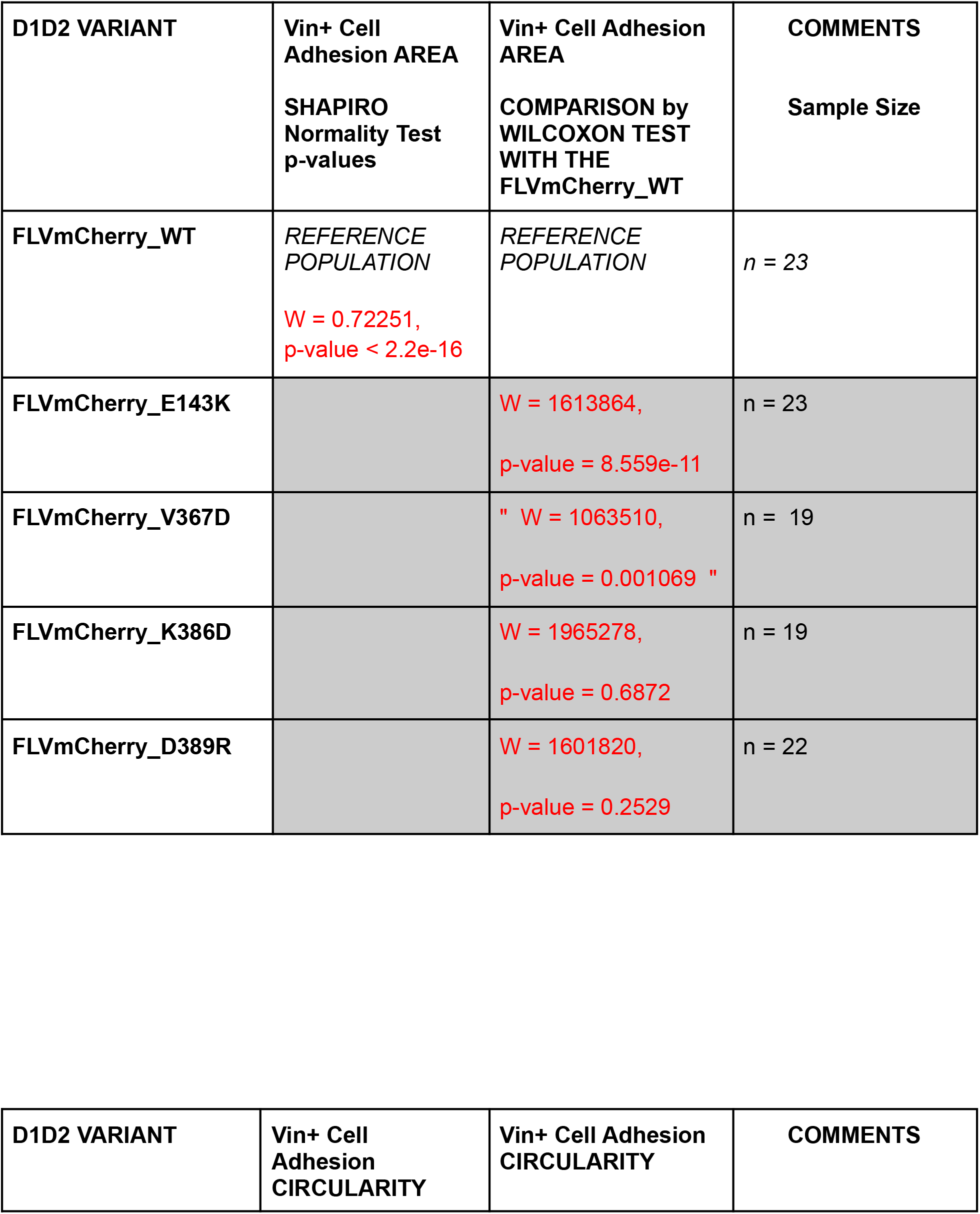
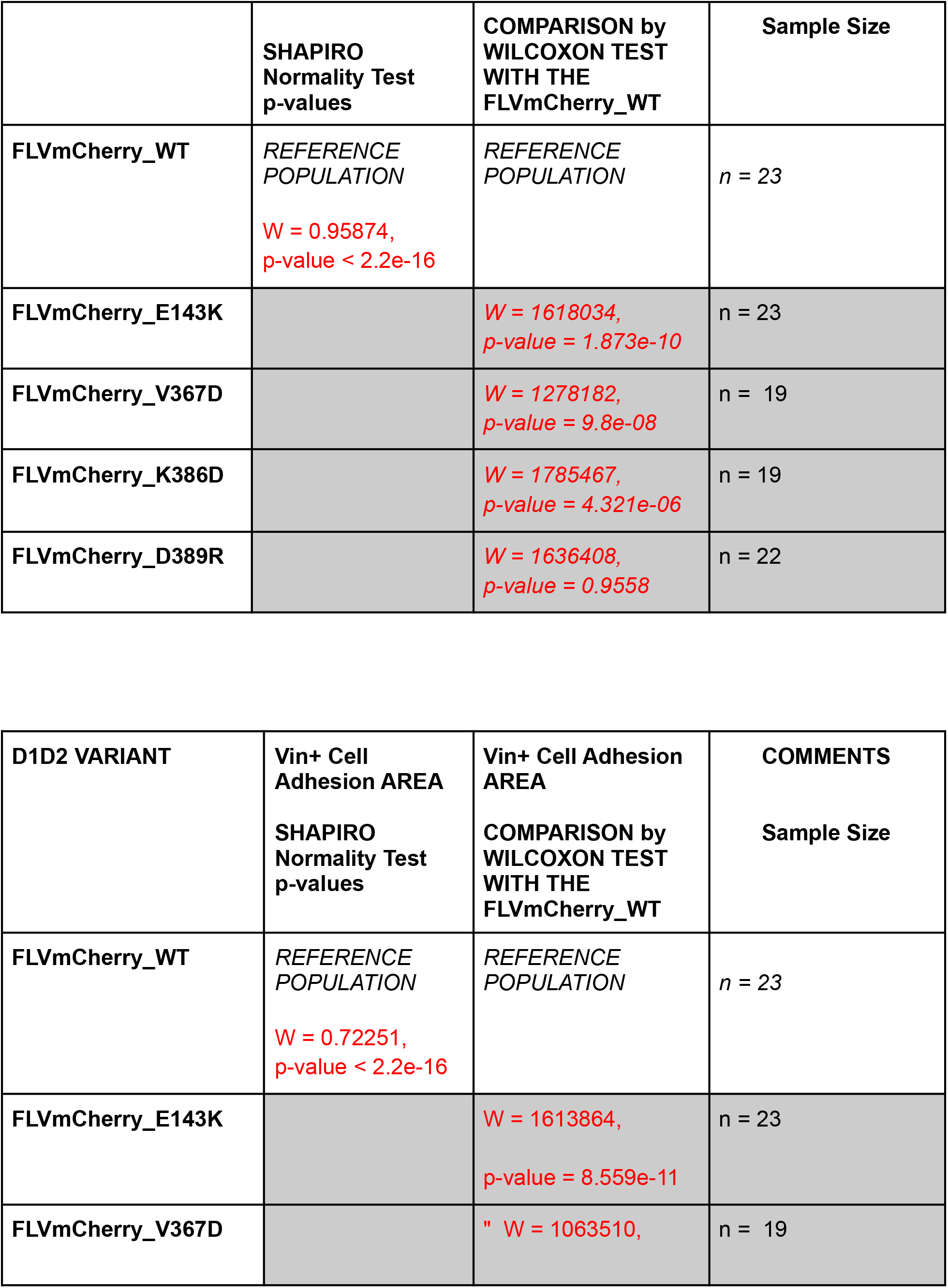
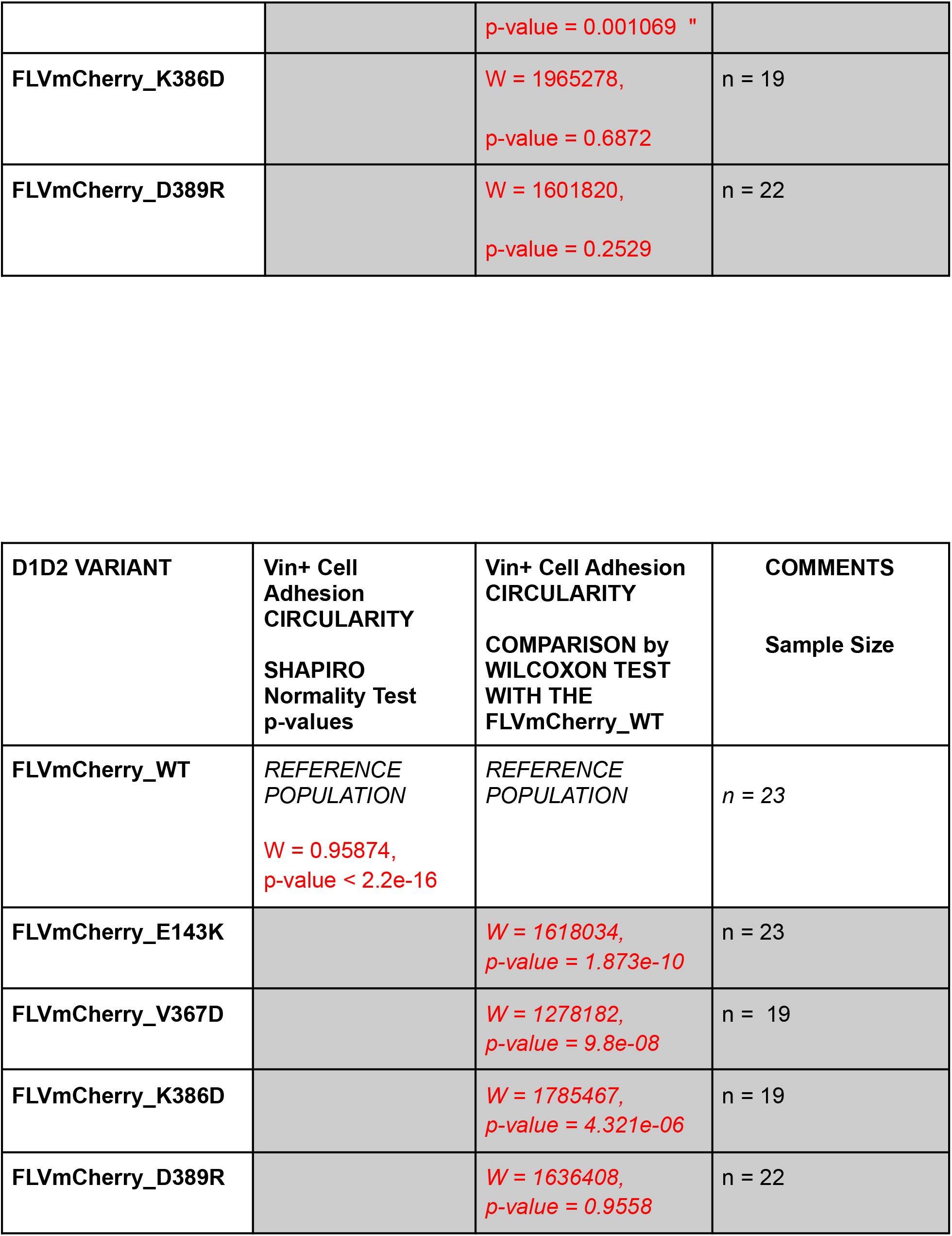
Values for the statistical analysis not included in the cell transfection experiments.

| D1D2 VARIANT | Vin+ Cell Adhesion AREA<br>SHAPIRO Normality Test p-values | Vin+ Cell Adhesion AREA<br>COMPARISON by WILCOXON TEST WITH THE FLVmCherry_WT | COMMENTS<br>Sample Size |
| --- | --- | --- | --- |
| FLVmCherry_WT | REFERENCE POPULATION<br>W = 0.72251,<br>p-value < 2.2e-16 | REFERENCE POPULATION | n = 23 |
| FLVmCherry_E143K |  | W = 1613864,<br>p-value = 8.559e-11 | n = 23 |
| FLVmCherry_V367D |  | " W = 1063510,<br>p-value = 0.001069 " | n = 19 |
| FLVmCherry_K386D |  | W = 1965278,<br>p-value = 0.6872 | n = 19 |
| FLVmCherry_D389R |  | W = 1601820,<br>p-value = 0.2529 | n = 22 |

| D1D2 VARIANT | Vin+ Cell Adhesion CIRCULARITY | Vin+ Cell Adhesion CIRCULARITY | COMMENTS |
| --- | --- | --- | --- |

|  | **SHAPIRO**<br>**Normality Test**<br>**p-values** | **COMPARISON by**<br>**WILCOXON TEST**<br>**WITH THE**<br>**FLVmCherry_WT** | **Sample Size** |
| --- | --- | --- | --- |
| FLVmCherry_WT | *REFERENCE*<br>*POPULATION*<br>*W = 0.95874,*<br>*p-value < 2.2e-16* | *REFERENCE*<br>*POPULATION* | *n = 23* |
| FLVmCherry_E143K |  | *W = 1618034,*<br>*p-value = 1.873e-10* | n = 23 |
| FLVmCherry_V367D |  | *W = 1278182,*<br>*p-value = 9.8e-08* | n = 19 |
| FLVmCherry_K386D |  | *W = 1785467,*<br>*p-value = 4.321e-06* | n = 19 |
| FLVmCherry_D389R |  | *W = 1636408,*<br>*p-value = 0.9558* | n = 22 |

| **D1D2 VARIANT** | **Vin+ Cell**<br>**Adhesion AREA**<br>**SHAPIRO**<br>**Normality Test**<br>**p-values** | **Vin+ Cell Adhesion**<br>**AREA**<br>**COMPARISON by**<br>**WILCOXON TEST**<br>**WITH THE**<br>**FLVmCherry_WT** | **COMMENTS**<br>**Sample Size** |
| --- | --- | --- | --- |
| FLVmCherry_WT | *REFERENCE*<br>*POPULATION*<br>*W = 0.72251,*<br>*p-value < 2.2e-16* | *REFERENCE*<br>*POPULATION* | *n = 23* |
| FLVmCherry_E143K |  | *W = 1613864,*<br>*p-value = 8.559e-11* | n = 23 |
| FLVmCherry_V367D |  | *" W = 1063510,* p-value = 0.001069 " | n = 19 |
| FLVmCherry_K386D |  | W = 1965278,<br>p-value = 0.6872 | n = 19 |
| FLVmCherry_D389R |  | W = 1601820,<br>p-value = 0.2529 | n = 22 |

| D1D2 VARIANT | Vin+ Cell Adhesion CIRCULARITY<br>SHAPIRO Normality Test p-values | Vin+ Cell Adhesion CIRCULARITY<br>COMPARISON by WILCOXON TEST WITH THE FLVmCherry_WT | COMMENTS<br>Sample Size |
| --- | --- | --- | --- |
| FLVmCherry_WT | REFERENCE POPULATION<br>W = 0.95874,<br>p-value < 2.2e-16 | REFERENCE POPULATION | n = 23 |
| FLVmCherry_E143K |  | W = 1618034,<br>p-value = 1.873e-10 | n = 23 |
| FLVmCherry_V367D |  | W = 1278182,<br>p-value = 9.8e-08 | n = 19 |
| FLVmCherry_K386D |  | W = 1785467,<br>p-value = 4.321e-06 | n = 19 |
| FLVmCherry_D389R |  | W = 1636408,<br>p-value = 0.9558 | n = 22 |

## Notes

### Competing Interest Statement

The authors have declared no competing interest.

